# Ankyrin B Promotes Developmental Spine Regulation in the Mouse Prefrontal Cortex

**DOI:** 10.1101/2023.07.11.548527

**Authors:** Kelsey E. Murphy, Bryce W. Duncan, Justin E. Sperringer, Erin Y. Zhang, Victoria A. Haberman, Elliott V. Wyatt, Patricia F. Maness

## Abstract

Postnatal regulation of dendritic spine formation and refinement in cortical pyramidal neurons is critical for excitatory/inhibitory balance in neocortical networks. Recent studies have identified a selective spine pruning mechanism in the mouse prefrontal cortex (PFC) mediated by class 3 Semaphorins and the L1-CAM cell adhesion molecules Neuron-glia related CAM (NrCAM), Close Homolog of L1 (CHL1), and L1. L1-CAMs bind Ankyrin B (AnkB), an actin-spectrin adaptor encoded by *Ankyrin2* (*ANK2*), a high confidence gene for autism spectrum disorder (ASD). In a new inducible mouse model (Nex1Cre-ERT2: *Ank2*^flox^: RCE), *Ank2* deletion in early postnatal pyramidal neurons increased spine density on apical dendrites in PFC layer 2/3 of homozygous and heterozygous *Ank2*-deficient mice. In contrast, *Ank2* deletion in adulthood had no effect on spine density. Sema3F-induced spine pruning was impaired in cortical neuron cultures from AnkB-null mice and was rescued by re-expression of the 220 kDa AnkB isoform but not 440 kDa AnkB. AnkB bound to NrCAM at a cytoplasmic domain motif (FIGQY^1231^), and mutation to FIGQH inhibited binding, impairing Sema3F-induced spine pruning in neuronal cultures. Identification of a novel function for AnkB in dendritic spine regulation provides insight into cortical circuit development, as well as potential molecular deficiencies in ASD.

## Introduction

Developmental regulation of dendritic spine formation and remodeling in pyramidal neurons is necessary to achieve excitatory/inhibitory balance in neocortical circuits. During early stages of development, dendritic spines and excitatory synapses on cortical pyramidal neurons are initially overproduced, then selectively remodeled by elimination mechanisms that are incompletely understood (Huttenlocher PR 1979; Alvarez VA *et al*. 2007; Petanjek Z *et al*. 2011). In the prefrontal cortex (PFC) appropriately balanced circuits are important for such vital functions as working memory, cognitive flexibility, and sociability. In Autism Spectrum Disorders (ASD) these behaviors are impaired, and spine density is elevated in frontal cortical areas, which could contribute to network hyper-excitability (Hutsler JJ *et al*. 2010; Tang G *et al*. 2014; Forrest MP *et al*. 2018).

Recent investigation into molecular mechanisms of spine elimination in the mouse neocortex demonstrated that secreted Semaphorins of class 3 (Sema3) prune distinct populations of dendritic spines during development and homeostatic scaling (Tran TS *et al*. 2009; Wang Q *et al*. 2017; Mohan V, CS Sullivan*, et al.* 2019; Mohan V, SD Wade*, et al.* 2019; Duncan BW, KE Murphy*, et al.* 2021). As shown in Figure 1A, Sema3 dimers bind to heterotrimeric receptors comprising L1 cell adhesion molecules (L1-CAMs), Neuropilins (Npn1/2), and PlexinAs (PlexA1-4), to activate intracellular signaling resulting in spine pruning (Duncan BW, V Mohan*, et al.* 2021). Neuron-glial related CAM (NrCAM), Npn2, and PlexA3 constitute a receptor complex for Sema3F (Demyanenko GP *et al*. 2014; Mohan V, CS Sullivan*, et al.* 2019), while Close Homolog of L1 (CHL1), Npn2, and PlexA4 form a receptor complex for Sema3B (Mohan V, SD Wade*, et al.* 2019). Mouse genetic knockouts of NrCAM (Demyanenko GP *et al*. 2014), CHL1 (Mohan V, SD Wade*, et al.* 2019) or L1 (Murphy KE, SD Wade*, et al.* 2023) increased density of immature spines on apical dendrites of cortical pyramidal neurons. Sema3F and Sema3B are secreted by neurons in an activity-dependent manner, consistent with the idea that less active or immature spines may be pruned by Sema3s released from active synaptic neighbors to refine cortical circuits in development (Wang Q *et al*. 2017; Mohan V, CS Sullivan*, et al.* 2019).

**Figure 1:**
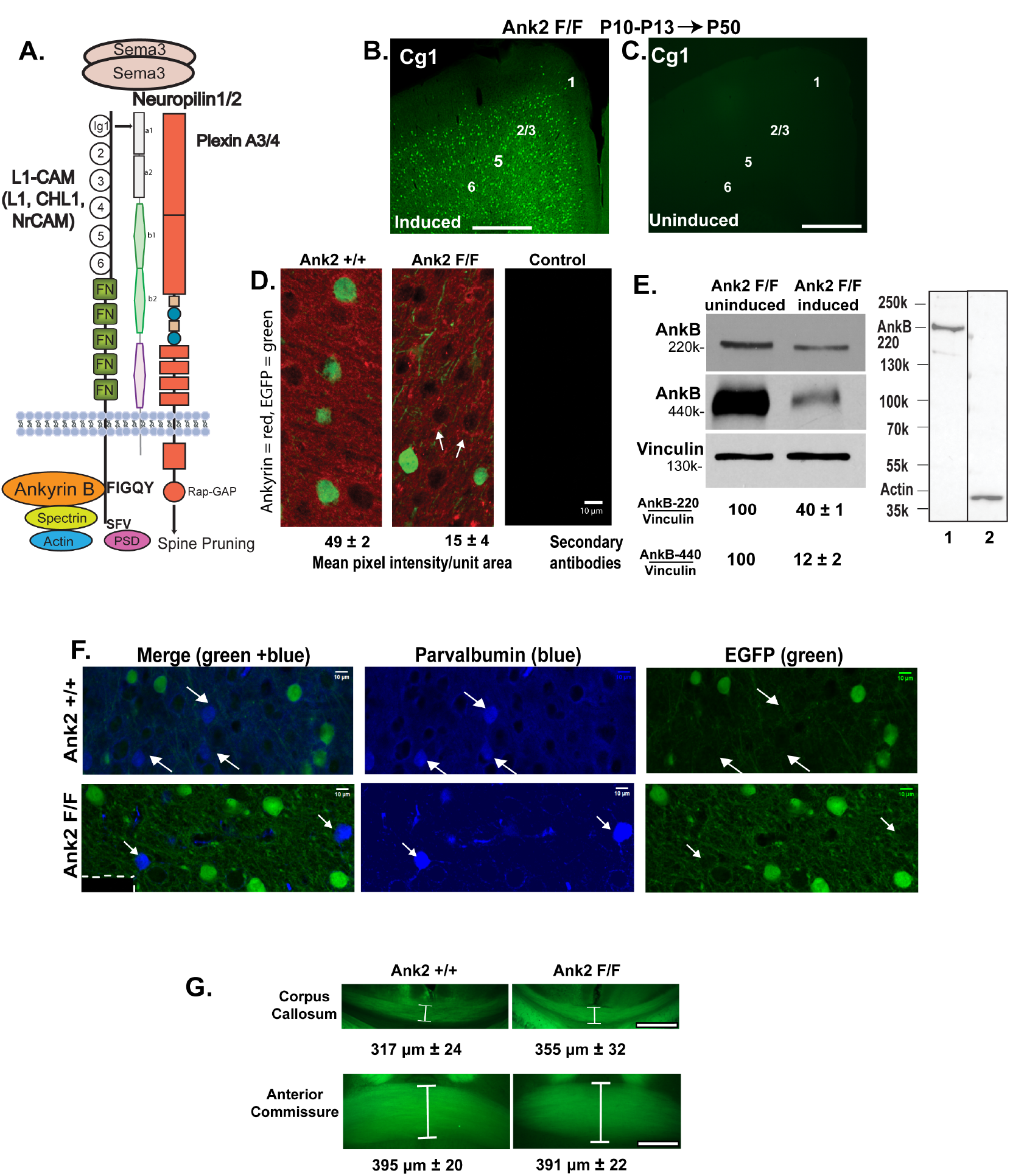
Conditional Deletion of AnkB in Postnatal Pyramidal Neurons of Nex1Cre-ERT2: *Ank2^F/F^*: RCE Mice. (A) Scheme of secreted class 3 Semaphorin dimers (Sema3) binding to heterotrimeric receptors at the neuronal membrane consisting of L1 cell adhesion molecules (L1-CAMs), Neuropilins (Npn1/2), and PlexinAs (PlexA1-4). This binding activates intracellular signaling through PlexA Ras-GAP activity, which induces spine pruning. Cytoplasmic domain motifs of L1-CAM bind Ankyrin B (FIGQY) and PSD proteins (SFV). Ankyrin B engages Spectrin, which links to the submembrane actin cytoskeleton. (B) Recombination in *Ank2* F/F primary cingulate cortex (Cg1) at P50 indicated by EGFP immunofluorescence (Alexafluor-488) in pyramidal neurons after TMX induction (P10-13). Cortical layers 1-3, 5, and 6 are labeled. Bar = 100 µm. (C) EGFP-labeled neurons are not evident in Cg1 of uninduced *Ank2* F/F mice (P50). Bar = 100 µm. (D) Representative confocal images showing EGFP-labeled pyramidal neurons (green, Alexafluor-488) and reduced immunofluorescence of AnkB protein labeled with AnkB monoclonal antibody (Thermofisher, 33-3700) (red, Alexafluor-555) in Cg1 layer 2/3 following TMX induction of *Ank2* F/F mutant mice compared to *Ank2* +/+ mice. Single optical sections were captured at the same confocal settings. Mean pixel intensity (± SEM) of AnkB immunofluorescence/unit area (below) was significantly lower in *Ank2* F/F compared to *Ank2* +/+ mice (n= 3 mice/genotype; 7 images/mouse; Mann-Whitney 2-tailed test, p=0.03). Arrows in *Ank2* F/F cortex point to residual AnkB immunofluorescence at the perisomatic region of EGFP-nonexpressing cells. Negative control with both AlexaFluor-conjugated secondary antibodies without primary antibodies showed no detectable fluorescence. All images were captured at same confocal settings. (E) Representative AnkB immunoblots of cortical lysates (5 µg) from uninduced and induced *Ank2* F/F mice (P50, n= 3 mice/condition). Densitometric scanning of protein bands from multiple blots showed that the ratios of AnkB-220 (antibody 33-3700) or AnkB-440 (sheep polyclonal antibody) relative to vinculin as control decreased significantly following TMX induction in *Ank2* F/F cortex compared to uninduced *Ank2* F/F cortex (mean ratio ± SEM; Mann-Whitney 2-tailed test, p=0.01 for each isoform). Ratios in uninduced cortex were set at 100. Different SDS-PAGE and transfer conditions were required to resolve AnkB-220 and AnkB-440 isoforms. (Right): Immunoblotting of cortical lysates under optimal conditions for AnkB-220 shows the specificity of AnkB antibody 33-3700 used for immunostaining, and that AnkB is not recognized by an irrelevant control antibody, anti-actin. (F) *Ank2* +/+ and *Ank2* F/F mice were induced with TMX and immunostained at P50 for parvalbumin (blue) and EGFP (green). Single channels show immunostaining of parvalbumin (blue channel) and EGFP (green channel), with merged images in first panels. Insert in lower first panel (dotted line) shows control staining with secondary AlexaFluor antibodies alone. Parvalbumin+ basket interneurons (arrows) are EGFP-negative. Images are maximum intensity confocal projections (scale bar = 10 μm). (G) Representative confocal images of the corpus callosum and anterior commissure at the midline (white brackets) in coronal brain sections of *Ank2* +/+ mice (n=4) and *Ank2* F/F mice (n=3, P50) expressing EGFP after TMX induction (P10-P13). Measurement of mean width (± SEM) were made using the straight-line tool in FIJI. Results indicated no significant difference in mean width of either tract for *Ank2* F/F compared to *Ank2* +/+ mice at the corpus callosum (Mann-Whitney test, 2-tailed, p= 0.39; scale bar = 50 µm) or anterior commissure (p=0.72; scale bar = 25 µm).

All L1-CAMs bind the actin-spectrin adaptor protein Ankyrin B (AnkB) at a conserved cytoplasmic domain motif (FIGQY) (Bennett V *et al*. 2009). A point mutation in the L1 Ankyrin binding site (FIGQH) is a pathological variant in the L1 intellectual disability syndrome with hydrocephalus (Hortsch M *et al*. 2014). L1 knock-in mouse mutants harboring this mutation display increased spine density in the prefrontal cortex (Murphy KE, SD Wade*, et al.* 2023) suggesting that AnkB may be vital for spine pruning *in vivo*. However, a recent study shows that this motif also binds Doublecortin-like kinase-1, a microtubule binding protein (Murphy KE, EY Zhang*, et al.* 2023).

Multiple *de novo* ASD variants in *ANK2* have been identified in the Simon Simplex Collection and Autism Sequencing Consortium (Iossifov I *et al*. 2012; Willsey AJ *et al*. 2013; De Rubeis S *et al*. 2014; Iossifov I *et al*. 2015; Wang T *et al*. 2016; Satterstrom FK *et al*. 2019; Wang T *et al*. 2020; Zhou X *et al*. 2022). *ANK2* is subject to alternative splicing resulting in a number of AnkB isoforms including a major isoform of 220 kDa, expressed in brain and other tissues, and a neuron-specific 440 kDa isoform (Jenkins PM *et al*. 2015). Over half of the 23 distinct variants in *ANK2* associated with ASD are predicted to be loss-of-function mutations affecting both principal AnkB isoforms.

Knowledge of AnkB function in developing brain is limited, because germline deletion of AnkB in mouse models is lethal in early postnatal life (Scotland P *et al*. 1998). To interrogate a potential role for AnkB in developmental spine regulation and to provide a model for loss-of-function ASD mutations, we generated a tamoxifen-inducible *Ank2* mouse line (Nex1Cre-ERT2: *Ank2^flox^*: RCE), which deletes all AnkB isoforms from postmitotic pyramidal neurons postnatally, circumventing lethality associated with germline deletion. Here we show that homozygous *Ank2* deletion in Nex1Cre-ERT2: *Ank2^flox^*: RCE mice during active stages of spine remodeling increased spine density and excitatory synaptic puncta on apical dendrites of prefrontal cortical pyramidal neurons, increasing the proportion of immature spines, whereas *Ank2* deletion in adulthood had no effect. Heterozygous mutants displayed an intermediate spine density phenotype. In wild type (WT) cortical pyramidal cells AnkB was localized to spines and dendrites and was enriched in mouse brain synaptoneurosomes. Cortical neurons in cultures from AnkB-null or NrCAM-null mice were deficient in Sema3F-induced spine pruning and specific re-expression of AnkB-220 but not AnkB-440 in *Ank2*-null neurons restored spine pruning. Mutation of tyrosine to histidine in the NrCAM FIGQY motif impaired both Sema3F-induced spine pruning and association with AnkB-220. These results suggest that AnkB loss or haploinsufficiency could alter developmental spine regulation and have consequences on cortical circuits relevant to ASD.

## Materials and Methods

### Generation and Tamoxifen Induction of Ank2 Conditional Mutant Mice

We generated an inducible *Ank2* mouse line Nex1Cre-ERT2: *Ank2^flox^*: EGFP^flox^ (C57Bl/6) to delete *Ank2* from post-mitotic, postmigratory cortical pyramidal neurons. Nex1Cre-ERT2 mice express a modified Cre recombinase fused to the human estrogen receptor, which is under control of endogenous Nex1 regulatory sequences and activated by tamoxifen (TMX) (Agarwal A *et al*. 2012). The Nex1 promoter drives the TMX-inducible Cre-ERT2 recombinase only in postmitotic pyramidal neurons and not interneurons, oligodendroglia, astrocytes, or non-neural cells (Agarwal A *et al*. 2012). Nex1-Cre mice have been extensively employed to target postmitotic pyramidal neurons with high specificity (Golonzhka O *et al*. 2015; Fame RM *et al*. 2016; Mohan V, CS Sullivan*, et al.* 2019; Miller DS *et al*. 2021). For our approach Nex1Cre-ERT2 mice were intercrossed with RCE: loxP reporter mice (from Gordon Fishell), in which induction of Cre-ERT2 recombines a “floxed stop cassette” enabling expression of EGFP in Cre-induced neurons (Sousa VH *et al*. 2009). The resulting mice were intercrossed with *Ank2^flox^* mice (from Peter Mohler), in which recombination deletes exon 24, which has 73 bp of coding sequence, resulting in a frameshift that places a stop codon in exon 25 (Smith SA *et al*. 2015; Roberts JD *et al*. 2019).

For *in vivo* induction, TMX (Sigma-Aldrich, #10540-29-1) was dissolved (10 mg/ml) in sunflower seed oil and administered by intraperitoneal injection at 100 mg/kg body weight every 24 hours for 4 (P10-P13) or 10 (P50-P60) consecutive days. Mice were analyzed at P50 (young adult) or P80 (older adult), respectively. Postnatal TMX induction in Nex1-CreERT2 mice has been shown to achieve cell-specific targeting of postmitotic cortical and hippocampal pyramidal neurons (Agarwal A *et al*. 2012). For breeding Nex1Cre-ERT2^+/+^: *Ank2^flox/+^*: RCE^+/+^ mice were crossed with *Ank2^flox^* mice, yielding expected ratios of ∼50% *Ank2^flox^* homozygotes (Nex1Cre-ERT2^+/-^: *Ank2^flox/flox^*: RCE^+/-^) and 50% *Ank2^flox/+^*heterozygotes (Nex1Cre-ERT2^+/-^: *Ank2^flox/+^*: RCE^+/-^). Mice with wild type (WT) *Ank2* alleles (Nex1Cre-ERT2^+/-^: *Ank2^+/+^*: RCE^+/-^) were produced by intercrossing heterozygotes. Each allele was genotyped by PCR from genomic DNA.

Other mouse strains included wild type (WT) mice (JAX:000664) and *NrCAM*-null mice on the C57Bl/6 background (Sakurai T *et al*. 2001; Sakurai T *et al*. 2006). *Ank2*-null mice on a hybrid background Sv129/C57Bl were produced by intercrossing heterozygotes of each strain as described (Scotland P *et al*. 1998). Mice were maintained according to policies of the University of North Carolina Institutional Animal Care and Use Committee (IACUC; AAALAC Institutional Number: #329; ID# 18-073, 21-039) in accordance with NIH guidelines.

### Immunoreagents and Immunolabeling

Polyclonal antibodies were directed against the following proteins: NrCAM (Abcam #24344, RRID:AB_448024 or R&D Systems AF8538), Npn2 (R&D Systems, AF567), and GFP (Abcam #13970, RRID:AB_300798). Monoclonal antibodies were directed against AnkB (Thermo Fisher, #33-3700, RRID:AB_2533115; Biolegend, #821403, RRID:AB_2728536), vinculin (Thermofisher, #MA5-11690) and parvalbumin (Neuromab L11413). Sheep anti-AnkB antibodies were a gift from Paul Jenkins (University of Michigan) (Qu F *et al*. 2016). Also used were antibodies to vesicular glutamate transporter-1 (vGlut1, #821301, Biolegend) and Homer-1 (#160002, Synaptic Systems). Non-immune rabbit IgG (NIg), HRP-and AlexaFluor 488, 555, and 594-conjugated secondary antibodies were from Jackson Immunoresearch.

### Brain Fixation, Immunostaining, and Imaging

Mice at indicated postnatal ages were anesthetized with 2.5% Avertin, perfused transcardially with 4% paraformaldehyde (PFA)/PBS and processed for staining as described (Demyanenko GP *et al*. 1999). Brains were postfixed in 4% PFA overnight at 4°C, followed by 0.02% PBS-azide, then sectioned coronally on a vibratome (60 µm) and mounted on glass slides. Sections were permeabilized in 0.3% Triton X-100 and blocked in 10% normal donkey serum in PBS for 3 hours at room temperature, then incubated with chicken anti-GFP antibody (1:250) and/or mouse monoclonal anti-AnkB antibody 33-3700 (Thermofisher) (1:250) for 48 hours at 4°C. After washing, sections were incubated with anti-chicken Alexa Fluor 488 and/or anti-mouse Alexa Fluor 555 secondary antibodies (1:250) for 2 hours before mounting with Prolong Glass (Thermofisher). Confocal z-stacks were obtained by imaging on a Zeiss LSM 700 microscope in the UNC Microscopy Services Laboratory with an EC Plan Neofluar 40x objective using a 1.3 N.A. oil lens (0.3 µm optical sections). Images were acquired using a pinhole size of 1 AU. Zoom was adjusted to obtain pixel sizes of 0.13-0.14 µm. For AnkB localization in immunostained brain sections, pixel intensities were quantified from confocal maximum intensity projections in FIJI. Regions of interest (ROIs) were selected using a grid, and mean grey values obtained using the FIJI analysis menu to yield pixel intensities per unit area. Pixel intensities of negative control staining with secondary antibodies alone were obtained similarly to determine background fluorescence, which was subtracted from mean pixel intensities of AnkB immunofluorescence. The resulting mean pixel intensities per unit area for each genotype were compared by the Mann-Whitney test (two tailed, unequal variance) for significant differences (p<0.05). AnkB localization in cortical neurons in culture (DIV14) was analyzed by indirect immunofluorescence staining using sheep anti-AnkB (Qu F *et al*. 2016) and AlexaFluor secondary antibodies (Thermofisher #A-11039, RRID: AB-2534096). Stimulated emission depletion microscopy (STED) was used for AnkB localization in cortical neurons in culture by immunofluorescence staining with mouse monoclonal anti-AnkB antibody 33-3700 and AF-594-conjugated secondary antibodies (Abcam, #ab245992). STED images were acquired using a Leica SP8 STED confocal microscope with Lightning deconvolution in the UNC Neuroscience Research Center.

Excitatory synaptic puncta on spines of EGFP-expressing pyramidal neurons in Nex1Cre-ERT2:RCE and Nex1Cre-ERT2: *Ank2* F/F: RCE mice were analyzed confocally in coronal brain sections through the PFC in layer 2/3 after immunofluorescence staining of presynaptic vesicular glutamate transporter-1 (vGlut1) and postsynaptic Homer-1 with Alexafluor-conjugated secondary antibodies. Quantification of excitatory synaptic puncta was performed blind to observer in single optical confocal sections (0.3 µm) using Imaris software. The mean number of puncta per 10 µm dendrite length was calculated. Puncta densities were compared between genotypes by the Mann-Whitney test (two tailed, unequal variance; p < 0.05).

### Analysis of Dendritic Spine Density and Morphology

Spine densities of layer 2/3 pyramidal neurons in the PFC (primary cingulate area) were measured using Neurolucida software (MBF Bioscience) as described (Demyanenko GP *et al*. 2014; Mohan V, CS Sullivan*, et al.* 2019; Mohan V, SD Wade*, et al.* 2019). Briefly, EGFP-labeled spines were traced and quantified blind to the observer on 30 µm segments of the first branch of apical or basal dendrites from maximum intensity projections. Images were compared to confocal z-stacks to verify that spines emerged from dendritic segments. Mean spine number per 10 µm of dendritic length (density) was calculated. Neurons (n = 10-21) were analyzed for each genotype on apical or basal dendrites. For production of 3D reconstructions dendritic z-stacks were deconvolved using AutoQuant 3 software (Media Cybernetics) with default deconvolution settings in Imaris (Bitplane). Mean spine densities/10 μM dendrite length ± SEM were analyzed by 2-factor ANOVA with Tukey’s posthoc testing (p<0.05).

Once spines densities were determined, spine morphologies of the same layer 2/3 pyramidal neurons in the PFC (primary cingulate area, P50) were manually scored on constructed Z-projections as mushroom, stubby, or thin (filopodial) on apical dendrites. Spines were measured in three categories, spine length, head width, and neck width, using the straight-line tool in FIJI. Spine length was defined as the total length of the spine from its base at the dendrite to its terminus. Head width was defined as the width of a distinct head, or, when lacking a distinct head, the measurement of the widest point in the terminal third of the spine. Neck width was defined as the width of the distinct neck of the spine, or, when a distinct neck was not discernible, the widest point in the basal third of the spine. Based on these three measurements, mushroom spines were defined by a spine length >0.5 μm, and a head width to neck width ratio >1.5. Stubby spines were defined by a length <1.0 μm and a ratio <1.5, while thin spines were defined by a length >1.0 μm and a ratio >1.0. To compare spine morphologies between genotypes of mice in PFC (layer 2/3), spine morphological types were calculated as a percentage of the total spine number. Significant differences across genotypes were determined by P-values (<0.05) computed by one factor ANOVA with Tukey posthoc comparisons.

Dendritic arborization was measured by Sholl analysis of processes crossing concentric rings centered on the soma of EGFP-labeled pyramidal neurons in confocal z-stacks (20x). The center was defined as the middle of the cell body at a soma detector sensitivity of 1.5 µm, and the automatic tracing mode of Neurolucida was used to seed and trace dendritic arbors. Images in DAT format were subjected to Sholl analysis using Neurolucida Explorer (MBF Bioscience) with a starting radius of 10 µm and radius increments of 10 µm ending at 300 µm. Apical and basal dendrites were not able to be scored separately due to overlap at increasing distances from soma. Data were analyzed using two-factor ANOVA (p<0.05).

The midline widths of the corpus callosum and anterior commissure in TMX-induced Nex1Cre-ERT2:RCE and Nex1Cre-ERT2: *Ank2 ^flox^*: RCE mouse brains were measured in serial coronal brain sections (60 µm) after immunostaining with antibodies against GFP. Confocal images were captured and analyzed in FIJI. The dorsoventral width of each tract was measured at the midline of each section using the straight-line tool in FIJI, and mean widths calculated. Means (± SEM) were compared by the Mann-Whitney test (two-tailed, unequal variance).

### Cortical Neuron Cultures and Sema3F-Induced Spine Retraction

Cortical neurons were isolated from brains of WT, *NrCAM*-null, or *Ank2*-null embryos (E15.5) and plated onto Lab-Tek II chamber slides (1.5 x 10^6^ cells/well) coated with poly-D-lysine and laminin as described (Mohan V, CS Sullivan*, et al.* 2019). For *Ank2*-null primary cultures, rapid genotyping of embryos was done while cortices of embryos remained on ice in Hibernate-E media containing B27 Plus supplement. AraC was added at 5 days *in vitro* (DIV5) to limit the growth of glia and fibroblasts, and media was changed on DIV7. At DIV11, cells were transfected with pCAGG-IRES-mEGFP with or without additional plasmids (AnkB-220 (pEGFP-N1-ΔEGFP) or AnkB-440 (pAB)) using Lipofectamine 2000 (Mohan V, CS Sullivan*, et al.* 2019; Duncan BW, V Mohan*, et al.* 2021). At DIV 14, cultures were treated with purified Fc from human IgG (Abcam #ab90285) or recombinant mouse Sema3F-Fc fusion protein (R&D Systems #3237-S3) at 5 nM for 30 minutes. Cultures were fixed with 4% PFA, quenched with 0.1M glycine, permeabilized with 0.1% Triton X-100, and blocked with 10% donkey serum. Cells were incubated with chicken anti-GFP primary antibody and AlexaFluor AF488-conjugated goat anti-chicken secondary antibody (1:500), washed, and mounted. At least 10 images of apical dendrites of labeled pyramidal neurons were captured per condition. Apical dendrites were distinguished from basal dendrites by cell morphology. Pyramidal neurons extend a single prominent, thicker apical dendrite from the apex of the pyramidal-shaped soma, whereas they extend multiple thinner basal dendrites from the base of the soma (Spruston N 2008). Any ambiguous dendrites were not analyzed. Confocal z-stacks were obtained using 0.2 µm optical sections of field size 64.02 x 64.02 µm, using a 40x oil objective and 2.4x digital zoom, and subjected to deconvolution. Spines from maximum intensity projections were traced and scored using Neurolucida. Mean spine densities (number/10 µm ± SEM) were calculated and compared by two-factor ANOVA with Tukey’s posthoc testing (p<0.05).

### Immunoprecipitation and immunoblotting

Protein-protein interactions were assessed by co-immunoprecipitation and immunoblotted from cortical lysates of mouse forebrain, synaptoneurosomes, and transfected HEK293T cells. For preparation of mouse cortical lysates, forebrains were dissociated and subjected to Dounce homogenization for 20 strokes in RIPA buffer. Homogenates were incubated for 15 minutes on ice, then centrifuged at 16,000 x g for 10 minutes. The supernatant was retained, and protein concentration determined by BCA. Synaptoneurosomes were isolated as described (Villasana LE *et al*. 2006). Briefly, WT mice (P32) were anesthetized, decapitated and cortices were isolated. Following Dounce homogenization in Triton lysis buffer (20 mM Tris, pH 7.0, 150mM NaCl, 5 mM EDTA, 1 mM EGTA, 1% Triton X-100, 10 mM NaF, 1x Protease Inhibitor Cocktail), homogenates were sonicated, filtered, and centrifuged at 1000 x g for 10 minutes at 4°C. Pellets were resuspended in Triton lysis buffer, nutated, and centrifuged at 16,000 x g for 10 minutes at 4°C. Supernatants were retained as the synaptoneurosome fraction. Protein concentrations were determined by BCA. Postsynaptic density protein 95 (PSD95), AnkB, α-tubulin, and actin levels were quantified by western blotting in the synaptoneurosome fraction compared to the filtered homogenate and 1000xg supernatant fraction as described previously (Villasana LE *et al*. 2006; Demyanenko GP *et al*. 2014; Murphy KE, EY Zhang*, et al.* 2023). HEK293T cells were grown in DMEM/ gentamicin/kanamycin/10% FBS in a humidified incubator with 5% CO2. Cells were seeded at 2 × 10^6^ cells/100mm dish the day before transfection. Plasmids were transfected with Lipofectamine 2000 in Opti-MEM. Media was changed to DMEM after 18 h, and cells were lysed and collected 48 h post-transfection. Cells were harvested in Brij98 lysis buffer (1% Brij98, 10 mM Tris-Cl pH 7.0, 150 mM NaCl, 1mM EDTA, 1mM EGTA, 10 mM NaF, protease inhibitors (SigmaAldrich #P8340).

For immunoprecipitation, lysates of mouse forebrain (1 mg) or HEK293T cells (0.5 mg) were precleared for 30 minutes at 4°C using Protein A/G Sepharose beads (ThermoFisher). Precleared lysates (equal amounts of protein) were incubated with 3 µg rabbit polyclonal antibody to NrCAM (Abcam #24344) or nonimmune IgG (nIg) for 2 hr on ice. Protein A/G Sepharose beads were added for an additional 30 min with nutation at 4°C before washing with RIPA or Triton lysis buffer (synaptoneurosomes). Beads were washed 4 times, then immunoprecipitated proteins were eluted from the beads by boiling in SDS-PAGE sample buffer. Samples (50 µg) were subjected to SDS-PAGE (6%) and transferred to nitrocellulose. Membranes were blocked in TBST containing 5% nonfat dried milk and incubated overnight with primary antibodies (1:1000), washed, and incubated with HRP-secondary antibodies (1:5000) for 1 h. Antibodies were diluted in 5% milk/Tris buffered saline/0.1% Tween-20 (TBST). For detection of AnkB-440, a 3.5-17.5% gradient gel was used, and AnkB antibodies were diluted in 5% BSA/TBST. Blots were developed using Western Bright ECL Substrate (Advansta) and exposed to film for times yielding a linear response of signal. Membranes were stripped and reprobed with rabbit anti-NrCAM antibodies (R&D Systems AF8538). Bands were quantified on densitometric scans in FIJI. The ratio of AnkB to NrCAM was determined from 3 replicate experiments to obtain mean and SEM values.

All experiments were designed to provide sufficient power (80–90%) to discriminate significant differences (p<0.05) in means (± SEM) between independent controls and experimental subjects as described (Dupont WD *et al*. 1990). The type I error probability associated with tests of the null hypothesis was set at 0.05. Full details of the experimental design, sample sizes, and number of repetitions for each experiment are reported in the figure legends.

## Results

### Postnatal deletion of Ankyrin B from prefrontal pyramidal neurons increases spine density and immature spine morphology

To identify consequences of postnatal AnkB deletion on dendritic spine remodeling in cortical pyramidal neurons, we generated a TMX-inducible mouse line, Nex1Cre-ERT2: *Ank2^flox^*: RCE (C57Bl/6). In Nex1Cre-ERT2:RCE mice, loxP recombination is activated by Cre-ERT2 recombinase under control of the Nex1 promoter in postmitotic pyramidal neurons treated with TMX (Agarwal A *et al*. 2012; Mohan V, CS Sullivan*, et al.* 2019). Reporter EGFP expression is induced by loxP recombination in the same cells (Sousa VH *et al*. 2009). To achieve *Ank2* deletion during the major postnatal period of spine formation and remodeling (Culotta L *et al*. 2020), mice were given daily TMX injections intraperitoneally from P10 to P13 as described (Agarwal A *et al*. 2012). During the P14-P21 time period dendritic spines are overproduced and pruned to appropriate levels, and spine remodeling decreases substantially from P19-P34 as mature circuits are established (Trachtenberg JT *et al*. 2002; Holtmaat AJ *et al*. 2005). *Ank2* deletion at these postnatal stages were chosen to overcome early postnatal lethality of germline *Ank2* deletion (Scotland P *et al*. 1998), thus circumventing effects on premigratory pyramidal cells and early neuronal precursors.

We focused on layer 2/3 pyramidal neurons of the PFC, because of their importance in social and cognitive circuits, which are affected in ASD (Yizhar O 2012; Kroon T *et al*. 2019). Treatment with TMX at P10-P13 efficiently upregulated EGFP expression in pyramidal neurons of young adult Nex1Cre-ERT2: *Ank2^flox/flox^*: RCE mice (termed Ank2 F/F) in the primary cingulate area (Cg1) of the PFC at P50 (Fig. 1 B, C). EGFP-expressing neurons were distributed across the cortical laminae in induced but not uninduced Ank2 F/F mice, indicating that Cre recombinase expression was tightly regulated. Prefrontal cortical regions in mice lack a canonical layer 4 (L4) as well as gene expression characterizing L4 neurons (Tasic B *et al*. 2016; Wang D *et al*. 2018; Anastasiades PG *et al*. 2021). Induced *Ank2* F/F mice were healthy and viable even as older adults (P150). AnkB immunofluorescence staining in PFC of TMX-induced Nex1Cre: *Ank2^+/+^*: RCE mice (termed *Ank2* +/+) was present in the neuropil, and appeared punctate, possibly reflecting spine and/or synaptic localization and appeared excluded from EGFP-negative nuclei or soma (Fig. 1D). AnkB immunolabeling was diminished in TMX-induced *Ank2* F/F mice compared to *Ank2* +/+, as shown in representative single optical sections captured at the same confocal settings (Fig. 1E). Negative control staining with secondary AlexaFluor secondary antibodies alone indicated that AnkB immunostaining in the *Ank2* F/F cortex was substantially greater than background (Fig. 1D, control). AnkB immunofluorescence was quantified by measuring pixel intensity per unit area in multiple single optical sections under the same confocal settings, and the resulting mean pixel intensity/unit area was significantly lower in *Ank2* F/F compared to *Ank2* +/+ PFC (Fig. 1D, below; Mann-Whitney, 2-tailed t-test, p = 0.03). Residual AnkB immunofluorescence in the *Ank2* F/F cortex was seen adjacent to EGFP-negative soma (arrows), and in the neuropil between soma (Fig. 1D). This immunoreactivity likely derived from interneuron processes, glia, or axons from neuronal cells whose soma are located in subcortical areas, such as the amygdala and ventral tegmental area, which do not express Nex1. In addition, pyramidal neurons with slightly different birthdates would be unaffected by TMX treatment at P10-P13. This was supported by expression of *Ank2* transcripts in pyramidal cells and interneurons, as well as in astrocytes and oligodendroglia of the adult mouse cortex (Allen Brain Map Transcriptomics Explorer (https://celltypes/brain-map.org/maseq).

Western blotting for AnkB in cortical lysates (P50) confirmed that there was decreased expression of both AnkB-220 and AnkB-440 following TMX induction in *Ank2* F/F forebrain compared to uninduced *Ank2* F/F (Fig. 1E). Densitometric measurements of AnkB from multiple blots showed that the mean density of each isoform decreased significantly upon induction relative to the vinculin control, but that AnkB-440 decreased to a greater extent than AnkB-220 (Fig. 1E, below; Mann-Whitney 2-tailed t-test, p=0.001 for each isoform). This was probably because neuron-specific AnkB-440 is under Nex1 control, whereas AnkB-220 expression in interneurons and glia is Nex1-independent. Note that optimal AnkB-220 and -440 detection on Western blots requires different electrophoresis and transfer conditions due to their large size and difference in molecular weight. Under conditions for Ank-220 the AnkB antibody (mouse monoclonal 33-3700) recognized a single AnkB-220 band in cortical lysates, while an irrelevant control antibody (anti-actin) did not recognize AnkB or any other protein (Fig. 1E, right).

Basket interneurons account for ∼ 50% of cortical interneurons, of which parvalbumin-expressing basket cells constitute the majority (Markram H *et al*. 2004). To validate the specificity of Nex1Cre-ERT2 induced recombination, we carried out double immunostaining for parvalbumin and EGFP in PFC layer 2/3 of TMX induced *Ank2* F/F and *Ank2* +/+ mice (P50) (Fig. 1F). Parvalbumin+ interneurons in *Ank2* +/+ and *Ank2* F/F cortex (blue, arrows) were distinct from EGFP-expressing pyramidal neurons (green). There was no major alteration in the size of the corpus callosum or anterior commissure, major tracts comprising contralaterally crossing pyramidal cell axons. The mean width of EGFP-labeled axonal tracts were not different between genotypes as measured in serial coronal sections of *Ank2* F/F and *Ank2* +/+ brains (Fig. 1G). Gross neuroanatomical defects including the size of brain ventricles (Iateral and third) and cortical thickness were also not observed in *Ank2* F/F mice (not shown).

To investigate an *in vivo* role of AnkB in dendritic spine regulation in the postnatally developing PFC and effect of gene dosage, recombination was induced in juvenile *Ank2* F/F, *Ank2* F/+, and *Ank2* +/+ mice at P10-P13 and analyzed in young adults at P50 (Laviola G *et al*. 2003). During this time frame dendritic spines are actively overproduced and spine turnover occurs concurrently, followed by a decrease in spine number from P19-P34 as mature circuits are stabilized (Trachtenberg JT *et al*. 2002; Holtmaat AJ *et al*. 2005). A significant increase in mean density (88%) of EGFP-labeled spines on apical dendrites of pyramidal neurons in PFC layer 2/3 was observed in homozygous *Ank2* F/F mice compared to *Ank2* +/+ mice (Fig. 2A, B; ANOVA with Tukey’s posthoc tests *p<0.0001). Homozygous *Ank2* mice reflect AnkB loss-of-function, whereas heterozygotes can reflect haplo-insufficiency associated with neurological diseases (Lim ET *et al*. 2013). Accordingly, we compared spine density in layer 2/3 pyramidal neurons of *Ank2* heterozygotes (P50) induced at P10-P13. *Ank2* heterozygotes displayed a significant increase (36.5%) in mean spine density on apical dendrites compared to *Ank2* +/+ mice (ANOVA with Tukey’s posthoc tests *p=0.02) and a significant decrease in spine density (27%) compared to *Ank2* F/F homozygotes (ANOVA with Tukey’s posthoc tests *p=0.001, Fig. 2A, B), indicative of an intermediate phenotype. In contrast spine densities on basal dendrites of layer 2/3 pyramidal cells were not significantly different across any of the genotypes (*Ank2* +/+, F/+, F/F mice; Fig. 2B). Because *Ank2* remains deleted after recombination induced by TMX treatment (P10-P13), effects seen at P50 could result from altered spine turnover from a few days after TMX injection until the time of analysis. As shown in mouse somatosensory cortex the fraction of spines lost by pyramidal neurons in L2/3 or L5 was greater than spines gained from P16-24, whereas at P28 rates were balanced and remained relatively stable (Trachtenberg JT *et al*. 2002; Holtmaat AJ *et al*. 2005). It is worth noting that the presence of spines in homozygous *Ank2*-deficient mice indicates that AnkB is not required during this period to maintain spine integrity. Spines may be stabilized by AnkG, a different gene product localized to adult spine nanodomains, where it promotes spine maintenance and plasticity (Smith KR *et al*. 2014; Yoon S *et al*. 2020).

**Figure 2:**
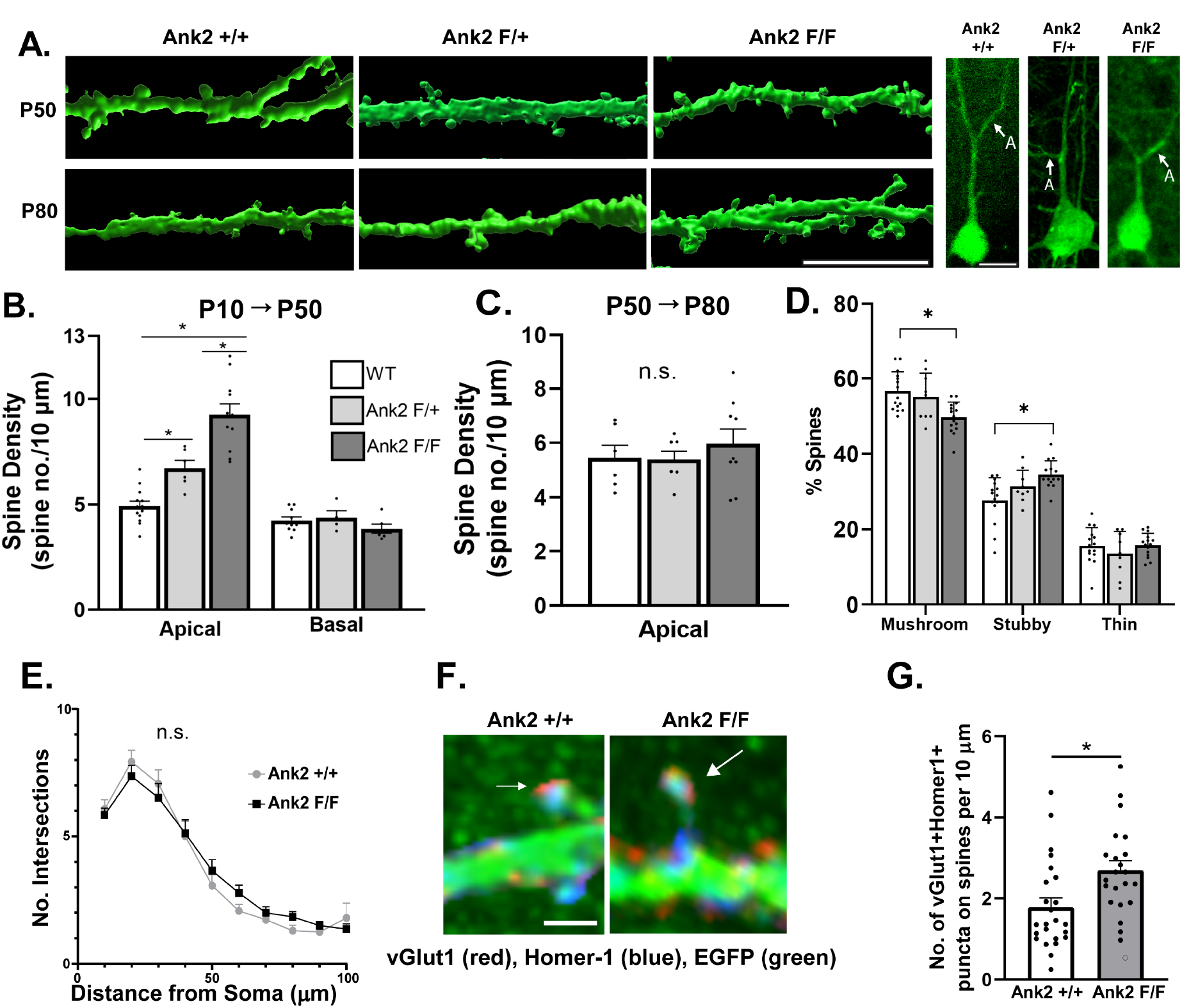
Increased Spine Density and Immature Synapses on Apical Dendrites of Ank*2* F/F Pyramidal Neurons in PFC Layer 2/3. (A) Representative images of EGFP-labeled apical dendrites and spines on pyramidal neurons in layer 2/3 of the medial PFC were subjected to 3D reconstruction of deconvolved dendritic z-stacks in Imaris (scale bar = 7 µm). Upper panels show that deletion of *Ank2* at P10-P13 by TMX induction increased spine density on apical dendrites in *Ank2* F/F and *Ank2* F/+ mice compared to *Ank2* +/+ mice at P50. Lower panels show that deletion of *Ank2* by TMX induction at P50 did not increase spine density on apical dendrites in older adults at P80. Three panels at right are representative low magnification confocal images of cortical neurons in PFC layer 2/3 from P50 mice, with arrows indicating apical dendrites (A) oriented toward the pial surface. Scale bar = 20 µm. (B) Quantification of mean spine density per 10 µm dendritic length (± SEM) on apical and basal dendrites of pyramidal neurons in layer 2/3 of PFC in *Ank2* +/+, F/+, and F/F mice induced at P10-P13 and analyzed at P50. Each point represents the mean spine intensity per mouse analyzed. One-factor ANOVA with Tukey’s posthoc testing showed significantly increased spine density on apical dendrites of *Ank2* F/F pyramidal neurons (9.2 spines/10 µm ± 0.5, n=11 mice) compared to *Ank2* +/+ (4.9 spines/10 µm ± 0.2, n=13 mice, *p<0.0001). *Ank2* F/+ mice exhibited intermediate spine density on apical dendrites (6.7 spines/10 µm ± 0.4, n=6 mice), which was significantly greater than *Ank2* +/+ mice (*p=0.02), and significantly less than *Ank2* F/F (*p=0.001). Spine density on basal dendrites of *Ank2* +/+ pyramidal neurons in layer 2/3 PFC (4.2 spines/10 µm ± 0.2; n=10 mice) was not significantly different from heterozygous *Ank2* F/+ (4.4 spines/10 µm ± 0.3; n=4 mice, p = 0.91) or *Ank2* F/F mice (3.8 ± 0.2; n=6 mice, p = 0.39). *Ank2* F/+ mice were not significantly different from *Ank2* F/F mice (p = 0.33). (C) Quantification of mean spine density per 10 µm dendritic length (± SEM) on apical dendrites of pyramidal neurons in layer 2/3 of PFC in *Ank2* +/+, F/+, and F/F mice induced at P50 and analyzed at P80. Each point represents the mean spine density per mouse. One-factor ANOVA with Tukey’s posthoc testing showed no difference in apical dendritic spine density in *Ank2* +/+ mice (5.4 ± 0.5, n=6 mice) compared to *Ank2* F/+ mice (5.4 ± 0.3, n=6 mice, p = 0.99) or *Ank2* F/F mice (6.0 spines/10 µm ± 0.5; n=9 mice, p = 0.72). *Ank2* F/+ mice were also not significantly different from *Ank2* F/F mice (p = 0.64). (D) Spine morphologies were quantified in *Ank2* +/+, *Ank2* F/+, and *Ank2* F/F mice induced at P10-P13 and analyzed at P50. Quantities of each morphological type are reported as percentages of total number of spines (± SEM). Spines were identified as one of three morphologies based on spine length and the ratio of head width to neck width. There was a significantly increased percentage of stubby spines in *Ank2* F/F (34% of total spines ± 0.9) compared to *Ank2* +/+ mice (28% ± 1.5, one-factor ANOVA with Tukey’s posthoc testing *p = 0.005). There was a significant decrease in mushroom spines in *Ank2* F/F (50% ± 1.1) compared to *Ank2* +/+ mice (57% ± 1.3, *p = 0.004), There was not a significant difference between *Ank2* F/+ stubby spines (31% ± 1.4) and either *Ank2* F/F stubby spines (p = 0.80) or *Ank2* +/+ stubby spines (p = 0.67). There was no significant difference between *Ank2* F/+ mushroom spines (55% ± 2.1) and *Ank2* F/F mushroom spines (p = 0.16) or *Ank2* +/+ mushroom spines (p = 0.99). The percent of thin spines was not significantly different between any genotype (p > 0.05). The total number of neurons analyzed per genotype was 9-15. The number of spines analyzed was for *Ank2* +/+ 441; *Ank2* F/+ 137; Ank2 F/F 634 (total =1212). (E) Sholl analysis of neuronal process branching of EGFP-expressing pyramidal neurons in PFC layer 2/3 showed no significant differences (n.s.) in the number of intersections of branches (primarily dendrites) with concentric circles at various distances from the soma of *Ank2* +/+ and *Ank2* F/F mice (P50) following TMX induction at P10-P13 (n =15 cells/genotype, 3 mice/genotype, two-factor ANOVA, p = 0.85). (F) Excitatory synaptic puncta were visualized by immunofluorescence staining of juxtaposed vGlut1 with Alexafluor-555 (red) and Homer-1 with Alexafluor-647(blue) in layer 2/3 of the PFC on apical dendritic spines of EGFP-expressing pyramidal neurons in *Ank2* +/+ and *Ank2* F/F mice (P50) after TMX induction at P10-P13. Representative confocal images (merged) show synaptic puncta of adjacent vGlut1 and Homer-1 labeling (arrows), scale bar = 1 µm. (G) Quantification of vGlut1/Homer-1 double-labeled puncta on spines per 10 µm dendritic length in single optical sections. *Ank2* F/F neurons exhibited an increased mean density of these puncta compared to *Ank2* +/+ neurons (Mann-Whitney 2-tailed test, *p=0.004; n= 4 mice/genotype, 22-24 neurons/genotype). Points represent puncta density per neuron.

To determine if deletion of *Ank2* in adulthood alters spine density, *Ank2* F/F, *Ank2* F/+, and *Ank2* +/+ mice were TMX-treated in young adults at P50-P60 and analyzed for spine density in older adults at P80. There were no significant differences in spine density on apical dendrites of layer 2/3 pyramidal neurons in *Ank2* +/+, *Ank2* F/+, *Ank2* F/F PFC at P80 (Fig. 2A, C). These results suggested that spine regulation in the adult PFC, which may in part reflect homeostatic changes, is not substantially altered by depletion of AnkB in cortical pyramidal neurons.

Dendritic spines acquire diverse morphologies (mushroom, stubby, thin), which have different functional properties and are classified based on size and shape (Peters A *et al*. 1970). Spine morphologies are dynamically interchangeable, comprising a continuum from immature stubby and thin spines, which have immature synaptic contacts and PSDs associated with juvenile plasticity (Bourne J *et al*. 2007), to mushroom spines with mature synapses (Bhatt DH *et al*. 2009; Holtmaat A *et al*. 2009; Berry KP *et al*. 2017). Several studies have shown that immature spine morphology is associated with Fragile X syndrome (Phillips M *et al*. 2015) and intellectual disability (Forrest MP *et al*. 2018). In *Ank2* F/F mice TMX-induced deletion at P10-13 increased the percent of stubby spines and decreased the percent of mushroom spines on apical dendrites of layer 2/3 pyramidal neurons in the PFC at P53 (Fig. 2D). The differences between *Ank2* +/+ and *Ank2* F/F were statistically significant (ANOVA with Tukey posthoc testing, *p < 0.01; p values in Fig. 2D legend). There were no significant differences in the percent of stubby or mushroom spines in heterozygous Ank2 F/+ mice compared to *Ank2* +/+ or *Ank2* F/F mice, although a slight trend to increased stubby spines was noted (Fig. 2D). The percent of thin spines was not significantly different between any genotype. Stubby spines are prevalent in early postnatal life and sparse in adulthood, thus are more apt to undergo loss with developmental age. This is consistent with the hypothesis that AnkB mediates elimination of immature spines during postnatal development. For example, AnkB binding to L1-CAMs may stabilize or cluster Sema3 receptor complexes on the neuronal surface enabling Sema3-induced pruning of immature, less active stubby spines. However, due to the dynamic transformation among spine morphologies it is also possible that AnkB could promote loss of other spine types. The increase in stubby spines of the AnkB mutant PFC suggests an overall decreased stabilization of spines and excitatory synapses, which might hinder aspects of circuit consolidation. Because the secreted Semaphorin Sema3A can induce basal dendritic branching in pyramidal neurons (Tran TS *et al*. 2009), dendritic arborization (primarily dendrites) was evaluated by Sholl analysis of PFC layer 2/3 pyramidal cells at P50 following homozygous *Ank2* deletion at P10-P13. No differences were observed in dendritic branching of EGF-labeled *Ank2* F/F neurons compared to *Ank2* +/+ cells (Fig. 2E). Taken together, these results support a role for AnkB in constraining spine density and eliminating a fraction of immature spines on apical dendrites of layer 2/3 pyramidal neurons during circuit refinement in the PFC.

Immature spines with stubby or thin morphology have been shown to contain functional excitatory synapses albeit with a higher NMDA to AMPA receptor ratio than mature spines (Berry KP *et al*. 2017). These spines tend to be highly dynamic and can be transient, allowing for rewiring of connections as circuits mature. However, some spines, usually of thin or filopodial morphology, lack functional synapses with pre-and postsynaptic specializations. Excitatory synapses on dendritic spines contain presynaptic vesicular glutamate transporter 1 (vGlut1) and postsynaptic scaffold protein Homer-1. To assess whether the observed increase in spine density of *Ank2* F/F pyramidal cells resulted in a corresponding increase in excitatory synapses, synaptic puncta were identified on spines by juxtaposed immunofluorescence staining of vGlut1 and Homer-1 on EGFP-expressing apical dendrites in PFC layer 2/3 in brain sections of *Ank2* F/F and *Ank2* +/+ mice (P50). Juxtaposed VGlut1 and Homer-1 immunofluorescence was observed on EGFP-labeled spines of both genotypes (Fig. 2F). Quantification of dual-labeled puncta on spines revealed a significant increase in *Ank2* F/F compared to *Ank2* +/+ neurons (Fig. 2G). These results were consistent with an increase in functional excitatory synapses, rather than orphan synapses lacking pre-or postsynaptic specializations.

### AnkyrinB-220 Mediates Sema3F-Induced Spine Pruning

To assess a potential role for AnkB in Sema3F-induced spine pruning, and to evaluate AnkB isoform specificity in the response, we developed a spine retraction assay using *Ank2*-null cortical neuron cultures in which all *Ank2* isoforms are deleted. Most *Ank2*-null mice on a C57Bl/6 background die shortly after birth (Scotland P *et al*. 1998), and cortical neuron cultures from these embryos (E15.5) degenerated at approximately DIV6, prior to significant spine elaboration. We found that *Ank2*-null mice on a first-generation hybrid background C57Bl/6/SV129 survived to at least P5, and cortical neuron cultures from *Ank2*-null embryos (E15.5) were viable through DIV14, when spines were abundant. Intercrossing *Ank2* heterozygotes from each background strain produced WT and *Ank2*-null hybrid embryos in the same litters. Embryos (E15.5) were subjected to rapid genotyping then cortical neuron cultures were plated. Cortical cultures from WT and *Ank2*-null embryos were transfected on DIV11 with reporter plasmid pCAG-IRES-EGFP, then treated with Sema3F-Fc or Fc (5 nM, 30 min) on DIV14. After fixation and immunostaining for EGFP, spine density was quantified on apical dendrites of EGFP-expressing neurons. WT and Ank2-null neurons in control cultures treated with Fc exhibited equivalent spine densities (Fig. 3A, B). Neurons in culture from either genotype would not be expected to show differences in spine densities as there is little if any accumulation of Sema3B or Sema3F in culture supernatants due to multiple media changes. In contrast secreted Sema3s are present *in vivo* in brain where they could act to constrain spine density in WT but not *Ank2*-deficient mice. Such differences are also observed for WT and NrCAM-null or CHL1-null neurons in culture vs. *in vivo* (Demyanenko GP *et al*. 2014; Mohan V, CS Sullivan*, et al.* 2019; Mohan V, SD Wade*, et al.* 2019). In striking contrast, Sema3F-Fc decreased spine density in WT but not Ank2-null neurons, indicating that AnkB was required for the Sema3F pruning response (Fig. 3A, B). Spine retraction in response to Sema3F-Fc in WT neurons was incomplete, since only the subpopulation of spines expressing NrCAM responds to Sema3F, while CHL1-expressing neurons respond to Sema3B (Mohan V, SD Wade*, et al.* 2019).

**Figure 3:**
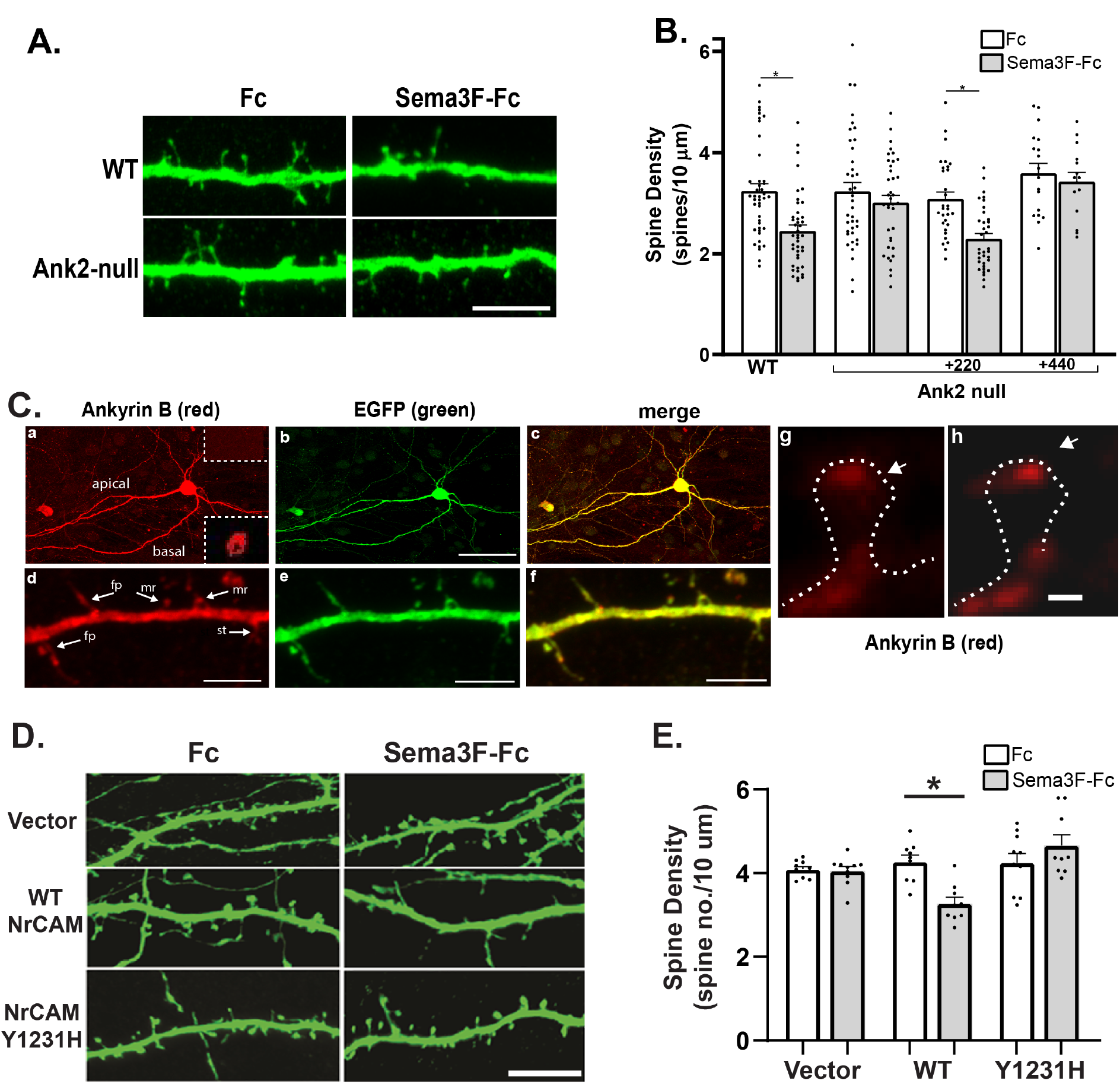
Sema3F-Induced Spine Retraction is Impaired in Cortical Neuron Cultures from *Ank2*-null, NrCAM-null, and NrCAM Y^1231^H Mutant Mice. (A) WT and *Ank2*-null cortical neuronal cultures were transfected with pCAG-IRES-EGFP, treated with 5 nM Fc or Sema3F-Fc for 30 min at DIV14, immunostained for EGFP, and apical dendrites imaged confocally (scale bar = 5 μm). *Ank2*-null cultures were co-transfected with Ank2-220 or Ank2-440 cDNAs. Representative images show EGFP-labeled apical dendrites with spines in WT and *Ank2*-null cortical neuronal cultures treated with Fc or Sema3F-Fc. Pial surface is toward the left. (B) Quantification of mean spine intensity ± SEM on apical dendrites of neuronal cultures described in (A). Each point represents the mean spine density per neuron per 10 µm dendrite length. Two-factor ANOVA with Tukey post hoc testing of spine density on *Ank2*-null neurons demonstrated that *Ank2* deletion inhibited Sema3F-induced spine retraction (Fc-treated, 3.23 spines/10 µm ± 0.18; Sema3F-Fc treated, 3.01 spines/10 µm ± 0.15, p = 0.99). In contrast WT neurons exhibited spine retraction in response to Sema3F (Fc-treated, 3.24 spines/10 µm ± 0.14; Sema3F-Fc-treated, 2.45 spines/10 µm ± 0.11, *p < 0.001). Sema3F-Fc-induced spine retraction in *Ank2*-null neurons was rescued by re-expressing AnkB-220 cDNA (Fc-treated, 3.08 spines/10 µm ± 0.14; Sema3F-Fc-treated, 2.30 spines/10 µm ±0.10, *p=0.001). AnkB-440 cDNA was not able to rescue Sema3F-induced spine retraction (Fc, treated, 3.59 spines/10 µm ± 0.20; Sema3F-Fc -treated, 3.43 spines/10 µm ±0.18, p = 0.99). Immunostaining with pan-AnkB antibodies verified equivalent levels of isoform expression (not shown). Points represent mean spine density of individual neurons. (C) WT cortical neuronal cultures were transfected with pCAG-IRES-EGFP, immunostained, and imaged confocally (DIV14). Representative confocal images of neurons with apical dendritic spines immunostained with pan-AnkB antibodies and Alexafluor-555 (red; a, d), GFP with Alexafluor-488 (green; b, e), and merged images (c, f). Control staining with secondary antibodies alone is shown in the upper right dotted inset in (a). Top panels are maximum intensity projections, with a single optical section of AnkB immunofluorescence staining of neurons shown in the lower right dotted inset in (a). AnkB immunofluorescence labeling was observed in spines with filopodial (fp), mushroom (mr), and stubby (st) morphology (d, arrows). At higher magnification AnkB immunolabeling (red) with Alexafluor-594 was evident within a spine head domain (arrows) by (g) confocal or (h) STED superresolution microscopy. Scale bars = 100 µm (a-c), 5 µm (d-f), 0.2 µm (g, h). (D) *NrCAM*-null cortical neuronal cultures were transfected with pCAG-IRES-EGFP vector alone, or plasmids containing WT or NrCAM Y^1231^H cDNAs. Cultures were treated with 5 nM Fc or Sema3F-Fc for 30 min at DIV14, immunostained for EGFP and apical dendrites imaged confocally (scale bar = 5 µm). Representative images of EGFP-labeled apical dendrites show that Sema3F-Fc promotes spine retraction only on *NrCAM*-null neurons re-expressing WT NrCAM. (E) Sema3F-Fc induces spine retraction on apical dendrites of *NrCAM*-null cortical neurons re-expressing WT NrCAM but not NrCAM Y^1231^H or empty vector. Two-factor ANOVA with Tukey post hoc testing showed a significant difference (*p=0.003) in spine density in neurons re-expressing WT NrCAM following treatment with control Fc (4.28 spines/10 µm ± 0.16) vs. Sema3F-Fc (3.28 spines/10 µm ± 0.15). In contrast there was not a significant difference (p = 0.47) in spine density in neurons re-expressing NrCAM Y^1231^H following treatment with control Fc (4.25 spines/10 µm ± 0.22) vs. Sema3F-Fc (4.67 spines/10 µm ± 0.25). There was also not a significant difference (p > 0.99) in spine density in NrCAM-null neurons transfected with vector alone.

The specific localization of AnkB-220, rather than AnkB-440, to the somato-dendritic domain of neurons (Stevens SR *et al*. 2021), led us to investigate whether AnkB-220 and/or -440 isoforms mediated Sema3F-induced spine retraction. Accordingly, the ability of each isoform to rescue spine pruning in AnkB-null neuronal cultures was assayed after transfection of the relevant cDNAs. Plasmids expressing AnkB-220 or AnkB-440 under control of the CMV enhancer/promoter were co-transfected on DIV11 into *Ank2*-null neurons with pCAG-IRES-EGFP, then treated on DIV14 with Sema3F-Fc or Fc. Expression of AnkB-220 effectively rescued Sema3F-induced spine retraction to the same extent as in WT neurons (Fig. 3B). In contrast, expression of AnkB-440 in AnkB-null neurons did not rescue Sema3F-induced spine retraction (Fig. 3B). These results supported the conclusion that AnkB-220 specifically mediates Sema3F-induced spine retraction *in vitro*.

To examine AnkB localization in neurons in culture, immunostaining was carried out on WT cortical neuron cultures (DIV14) transfected for EGFP expression. Immunolabeling with AnkB antibodies recognizing both isoforms (Qu F *et al*. 2016) showed AnkB localized to apical and basal dendrites as well as axons and soma in confocal images (Fig. 3C). AnkB immunostaining was evident on spines of mushroom (mr), filopodial (fp), and stubby (st) morphology (Fig. 3C, d arrows). Higher magnification confocal (g) and super-resolution STED microscopy (h) showed AnkB immunofluorescence in spine head domains (arrowheads) and adjacent dendritic segments (Fig. 3C). The presence of AnkB staining in diverse cellular compartments of neuronal cultures is consistent with previous work using pan-AnkB antibodies and AnkB-440 specific antibodies concluding that AnkB-220 preferentially localizes to the somato-dendritic domain of neurons (Stevens SR *et al*. 2021), whereas AnkB-440 is enriched in premyelinated axons (Kunimoto M *et al*. 1991; Kunimoto M 1995; Yang R *et al*. 2019). NrCAM is also present in all subcellular compartments of cortical neurons in culture (Demyanenko GP *et al*. 2014), although Npn2 is specifically localized to apical dendrites (Tran TS *et al*. 2009).

### The NrCAM Ankyrin Binding Motif is Required for Sema3F-Induced Spine Pruning

L1-CAMs bear a conserved cytoplasmic domain containing a FIG(Q/A)Y motif that reversibly binds Ankyrin (Bennett V *et al*. 2009). Phosphorylation of the tyrosine residue in the FIGQY motif can be achieved by receptor tyrosine kinases that are activated by NGF, bFGF (Garver TD *et al*. 1997) or EphrinB (Dai J *et al*. 2013), and this phosphorylation reverses Ankyrin binding to L1-CAMs (Bennett V *et al*. 2009). A charge reversal mutation of L1-FIGQY to FIGQH prevents Ankyrin association, and is a pathological mutation in the human L1 syndrome of intellectual disability (Hortsch M *et al*. 2014). In a knock-in mouse mutant expressing L1-FIGQH axonal connectivity is altered, as demonstrated by aberrant topographic mapping of retino-collicular axons (Buhusi M *et al*. 2008; Dai J *et al*. 2012). The L1-FIGQH mutation also suppresses axon branching (Yang R *et al*. 2019) and perisomatic innervation by interneurons (Tai Y *et al*. 2019). However, a role for Ankyrin association with L1-CAMs in spine elimination has not been demonstrated.

Structure-function studies showed that NrCAM binding to Npn2 increases binding between Npn2 and PlexA3, necessary for Sema3F-induced spine pruning (Duncan BW, V Mohan*, et al.* 2021). To investigate whether the Ankyrin binding motif FIGQY in the NrCAM cytoplasmic domain also promotes Sema3F-induced spine pruning, we analyzed the effect of the NrCAM FIGQY^1231^H mutation on Sema3F-induced spine retraction in neuronal cultures. NrCAM-null neurons in culture are refractory to Sema3F-induced spine retraction but re-expression of WT NrCAM rescues the response (Mohan V, CS Sullivan*, et al.* 2019). In the present study NrCAM-null neurons were transfected with pCAGS-IRES-EGFP plasmids encoding WT NrCAM or NrCAM-FIGQY^1231^H. Neuronal cultures were then treated with Sema3F-Fc or control Fc proteins (5 nM, 30 min) as described (Mohan V, CS Sullivan*, et al.* 2019). Spine density on apical dendrites of EGFP-expressing neurons was measured after immunostaining for EGFP. Sema3F-Fc promoted spine retraction in WT neurons but not in NrCAM-null cortical neurons, and re-expression of WT NrCAM restored the responsiveness to Sema3F-Fc, as reported (Mohan V, CS Sullivan*, et al.* 2019) (Fig. 3D, E). In contrast, NrCAM-null neurons transfected with the NrCAM FIGQY^1231^H mutant failed to respond to Sema3F-Fc and did not show a decrease in spine density (Fig. 3D, E). These results indicated that the Ankyrin binding site in the NrCAM cytoplasmic domain is a determinant of Sema3F-induced spine retraction.

### Ankyrin B Association with NrCAM

AnkB-220 and -440 isoforms are both expressed in mouse brain (Jenkins PM *et al*. 2015; Stevens SR *et al*. 2021), and share a membrane binding domain that binds L1-CAMs and certain ion channels (Chen K *et al*. 2017). Because of the selective ability of AnkB-220 to rescue Sema3F-induced spine pruning in AnkB-null cortical neurons, we examined the expression of AnkB-220 at different postnatal and adult stages in mouse cortex by Western blotting of forebrain lysates (equal protein). AnkB-220 was expressed at approximately equivalent levels in postnatal (P22, P34), young adult (P50) and older adult (P105) stages (Fig. 4A). NrCAM (130 kDa) also showed a relatively uniform expression pattern at these stages. To examine the association of AnkB-220 with NrCAM, NrCAM was immunoprecipitated from cortex lysates (P22-P105, equal protein), and immune complexes immunoblotted for AnkB-220 and NrCAM. AnkB-220 co-immunoprecipitated with NrCAM from cortex lysates at postnatal (P22, P34) and adult (P50, P105) stages (Fig. 4B). The relative amounts of associated AnkB-220 and NrCAM remained approximately equivalent from P22 to P50. The association of AnkB-220 with NrCAM was further assessed in synaptoneurosomes, a fraction enriched in pre-and postsynaptic terminals from mouse forebrain (P28) (Villasana LE *et al*. 2006). AnkB-220 co-immunoprecipitated with NrCAM from synaptoneurosome lysates (Fig. 4C). AnkB showed an enrichment in the synaptoneurosome fraction over filtered homogenate and S1 supernatants (equal protein), as did the postsynaptic density protein PSD95 (Fig. 4D), a known synaptic protein. Conversely, tubulin, a non-synaptic protein, had decreased expression in synaptoneurosomes when compared to filtered homogenate and S1 fractions (Fig. 4D) as noted previously (Villasana LE *et al*. 2006; Demyanenko GP *et al*. 2014).

**Figure 4:**
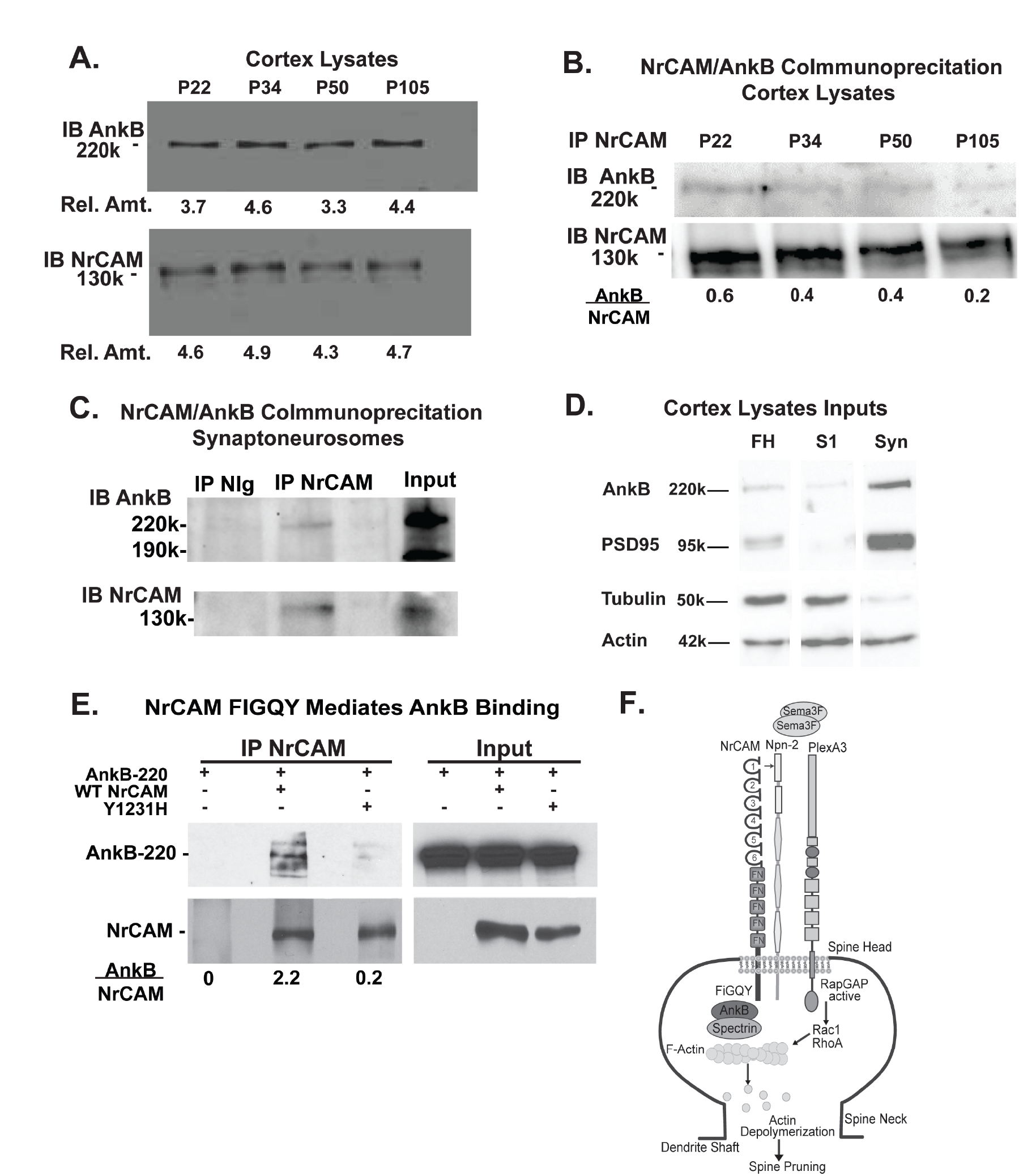
AnkB-220 Associates with NrCAM at the FIGQY Motif. (A) Representative immunoblotting (IB) of AnkB-220 (220kDa) and NrCAM (130 kDa) in cortical lysates from postnatal mouse brain (P22-P105) (20 µg protein). Relative amounts of AnkB-220 or NrCAM were determined by densitometric scanning of respective protein bands. AnkB-220 and NrCAM expression were equivalent from P22-P105. Similar results were obtained in duplicate experiments. (B) Co-immunoprecipitation of AnkB-220 (220kDa) and NrCAM (130 kDa) from equal amounts of protein (1mg) in cortex lysates of postnatal mouse forebrain (P22-P105). NrCAM was immunoprecipitated (IP) and AnkB-220 detected by immunoblotting (IB). AnkB immunoblots (above) were reprobed for NrCAM (below). Densitometric scanning was performed and the ratio of immunoprecipitated AnkB-220 to NrCAM indicated below. (C) Co-immunoprecipitation of AnkB-220 with NrCAM from P28 mouse synaptoneurosomes, shown by immunoprecipitation (IP) with NrCAM antibodies or nonimmune IgG (NIg) and immunoblotting (IB) with AnkB antibodies. Blots were reprobed by immunoblotting with NrCAM antibodies (lower panels). Inputs represent synaptoneurosome lysates that were not subjected to immunoprecipitation. Example is representative of replicate blots. (D) Filtered homogenate (FH), first supernatant (S1), and synaptoneurosome (Syn) samples (equal protein, 25μg) were blotted for Ankyrin B (AnkB). Membranes were stripped and reprobed for post-synaptic density protein 95 (PSD95, known synaptic protein), tubulin (a non-synaptic protein), and actin (loading control). AnkB and PSD95 showed enrichment in Syn fraction, whereas tubulin showed decreased expression in Syn fraction. (E) Co-immunoprecipitation of AnkB-220 with WT NrCAM or mutant NrCAM Y^1231^H from transfected HEK293T cells (equal amounts protein). NrCAM was immunoprecipitated from HEK293T cell lysates and immunoblotted for AnkB. Blots were reprobed for NrCAM (lower panels). Densitometric scanning of bands yielded ratios of AnkB-220 to NrCAM in the immunoprecipitated samples (below). Input lysates (equal protein) are shown at right. (F) Scheme of spine pruning initiated by Sema3F dimers. Sema3F binds the holoreceptor complex formed by NrCAM, Npn-2, and PlexA3. This binding event activates PlexinA3 Rap-GAP activity and subsequent downstream signaling leads to spine elimination via Rac1 and RhoA GTPase-governed pathways resulting in F-actin depolymerization. AnkB recruitment to the FIGQY motif in the NrCAM cytoplasmic domain may serve to stabilize the Sema3F complex, enhancing the signaling leading to spine pruning.

To probe the role of the NrCAM FIGQY motif in AnkB-220 binding, HEK293 cells were transfected with plasmids expressing AnkB-220 and either WT NrCAM or NrCAM FIGQY^1231^H. NrCAM was immunoprecipitated from equal amounts of HEK293 cell lysates then immunoblotted for AnkB. The relative amounts of AnkB/NrCAM in the immune complexes were quantified by densitometry. Results showed that AnkB-220 co-immunoprecipitated efficiently with WT NrCAM but much less with NrCAM FIGQY^1231^H (Fig. 4E). Input blots verified equivalent levels of AnkB-220, NrCAM, and NrCAM FIGQY^1231^H in cell lysates.

## Discussion

To investigate a role for AnkB in regulating dendritic spine regulation, we generated a conditional mouse model (Nex1Cre-ERT2: *Ank2^F/F^*: RCE) to inducibly delete *Ank2* from postmitotic, postmigratory pyramidal neurons at postnatal and adult stages in the PFC. Homozygous deletion of *Ank2* during early postnatal development when spines are most actively remodeled, resulted in elevated spine density on apical but not basal dendrites of layer 2/3 pyramidal neurons persisting in adulthood. Heterozygous deletion of *Ank2* postnatally increased spine density to an intermediate extent only on apical dendrites. In contrast, *Ank2* deletion in young adults did not alter spine density in older adult stages suggesting that mature functions of plasticity and homeostasis may be less impacted. Because apical dendrites have different synaptic inputs than basal dendrites (Brzdak P *et al*. 2019), it is possible that distinct synaptic connections are remodeled through AnkB interaction with L1 family adhesion molecules in response to different Semaphorins. However, the identity of such inputs has not been determined. L2/3 has an enormous variety of long-and short-range inputs and outputs. These include long range inputs from mediodorsal thalamus and basolateral amygdala, and short range outputs to L5 pyramidal tract cells, which target the thalamus and other sites to influence higher order behaviors of cognition, reward, and emotion (Anastasiades PG *et al*. 2021). Whereas we focused on AnkB, and its interaction with L1-CAMs, in the PFC, it is not regionally restricted to association cortical regions. For example, mouse null mutants in NrCAM and L1 genes cause increased spine density in pyramidal neurons in motor and visual cortex as well as in the PFC (Demyanenko GP *et al*. 2014; Murphy KE, SD Wade*, et al.* 2023).

AnkB was found to be essential for Sema3F-induced spine retraction, as demonstrated by a lack of response to Sema3F-Fc in *Ank2*-null neurons *in vitro*. Molecular replacement experiments in *Ank2*-null cortical neurons further revealed AnkB-220 as the isoform responsible for the Sema3F pruning response, rather than the neuron-specific isoform AnkB-440. This result was in accord with the preferential localization of AnkB-220 on neuronal dendrites and AnkB-440 on axons (Kunimoto M *et al*. 1991; Kunimoto M 1995; Chen K *et al*. 2020). Our localization studies, which were limited to the use of a pan-specific AnkB antibody, demonstrated the presence of AnkB within spines in addition to other compartments, but could not differentiate between AnkB-220 and AnkB-440.

The phenotype of elevated spine density on apical dendrites in Nex1Cre-ERT2: *Ank2^F/F^*: RCE mice is also present in *NrCAM* conditional mutant mice (Nex1Cre-ERT2: *NrCAM^F/F^*: RCE) (Mohan V, SD Wade*, et al.* 2019), as well as in Sema3F-null, Npn2-null, and PlexA3-null mice (Tran TS *et al*. 2009; Mohan V, SD Wade*, et al.* 2019). This elevated spine phenotype on apical dendrites is consistent with AnkB-mediated spine elimination through the Sema3F holoreceptor complex and was also observed in the PFC of CHL1-null and L1-null mutant mice (Murphy KE, SD Wade*, et al.* 2023). Since CHL1 shares a homologous Ankyrin binding site (FIGAY) and binds Npn2, Sema3B-induced retraction of a distinct spine subpopulation likely also depends on AnkB. Both Sema3F (Wang Q *et al*. 2017; Mohan V, CS Sullivan*, et al.* 2019) and Sema3B (Mohan V, SD Wade*, et al.* 2019) are expressed in an activity-dependent manner in culture, suggesting that these ligands may be released at active synaptic contacts to locally prune weak or inactive spines as cortical circuits mature. In line with this notion, we found that deletion of *Ank2* increased the relative proportion of stubby spines compared to mature mushroom spines, suggesting that immature spines may be selectively eliminated through AnkB.

NrCAM engaged AnkB-220 at a conserved FIGQY motif in the NrCAM cytoplasmic domain. This motif was required for spine elimination induced by Sema3F, as mutation of FIGQY to FIGQH disrupted AnkB-220 binding to NrCAM and blocked spine retraction in cortical neurons, analogous to a study in L1 knock-in mice with a substitution in the FIGQY motif (FIGQH) (Murphy KE, SD Wade*, et al.* 2023). As shown in the molecular scheme in Fig.4F, NrCAM binds Npn2 through extracellular domain interactions that increase Npn2/PlexA3 affinity (Mohan V, CS Sullivan*, et al.* 2019). Sema3F is a dimer that clusters and activates the holoreceptor complex, triggering the intrinsic Rap-GAP activity of PlexA3 (Wang Y *et al*. 2013). Downstream signal transduction ensues via pathways governed by Rac1 and RhoA GTPases leading to spine elimination on apical dendrites (Duncan BW, V Mohan*, et al.* 2021) (Fig. 4F). The recruitment of AnkB to the NrCAM cytoplasmic tail may stabilize the Sema3F holoreceptor complex, prolonging or enhancing signaling leading to spine retraction. Doublecortin-like kinase 1 (DCLK1) also binds to the Ankyrin binding motif FIGQY in NrCAM (Murphy KE, EY Zhang*, et al.* 2023). However, DCLK1 deletion in postnatal pyramidal neurons of Nex1Cre-ERT2: *DCLK1^flox/flox^*: RCE mice decreases spine density on apical dendrites in contrast to *Ank2* deletion, which increases spine density. Thus, NrCAM binding to Ankyrin, rather than DCLK1, most likely is responsible for constraining spine density.

The Nex1Cre-ERT2: *Ank2*: RCE mouse line reported here will be useful as a new animal model for *in vivo* studies relevant to ASD that involve prefrontal pyramidal circuits, such as social and cognitive behaviors. Our model most closely resembles ASD *de novo* truncating and frameshift (fs) mutations (for example, R895, R990, L1448, Q3683fs), in which both AnkB-220 and -440 isoforms are predicted to be deficient, and this model can be exploited to evaluate both homozygous and heterozygous deficiencies. Complete knockouts among all genes exome-wide significantly contribute to ASD (∼5% of cases) (Lim ET *et al*. 2013; Yu TW *et al*. 2013), thus it can be expected that there will be Ank2 homozygous or compound heterozygous variants effectively inactivating both alleles associated with ASD.

As shown here, *Ank2* F/F mice display phenotypes seen in ASD, including elevated spine density, and increased excitatory synaptic puncta. The line differs from a frameshift mouse mutant (ABe37fs/fs), in which a stop codon in exon 37 interrupts the neuron-specific sequence unique to AnkB-440 (Yang R *et al*. 2019). Juvenile ABe37fs/fs mice displayed transiently increased spines in the somatosensory cortex, and increased axonal branching in hippocampal cultures, however these effects were normalized by adulthood. Axons were not extensively analyzed in our *Ank2* F/F model but 2 major axon tracts, the corpus callosum and anterior commissure, were of normal size. Nex1Cre-induced deletion of *Ank2* at P10-P13 may minimize axon defects as deletion is restricted to postmitotic, postmigratory pyramidal neurons after many axons have reached their targets. Spine defects were not reported in a Nestin-Cre: *Ank2* exon37 F/F mutant in which 90% of AnkB-440 is lost from both neuronal and glial progenitors, or in a heterozygous truncation mutant (ABe22) targeting both AnkB-220 and -440 isoforms (homozygotes were lethal) (Yang R *et al*. 2019). All these mouse genetic models will be important in future studies to determine the relative roles of AnkB isoforms at different stages of dendritic and axonal development.

In conclusion, these results illuminate a new molecular function for AnkB in Sema3F-induced spine elimination in postnatally developing pyramidal neurons of the PFC. The findings contribute to our understanding of how cortical pyramidal cells may protect active spines while eliminating others in developing prefrontal circuits. This work also may provide molecular insight into ASD pathology associated with elevated spine density, hyper-excitability and over-connectivity in cortical circuitry (Hinton VJ *et al*. 1991; Irwin SA *et al*. 2001; Hutsler JJ *et al*. 2010; Tang G *et al*. 2014; Sellier C *et al*. 2017).

## Acknowledgements

We gratefully acknowledge Dr. Peter Mohler for providing Ank2 floxed mice, and Dr. Gordon Fishell for RCE mice. We acknowledge Dr. Paul Jenkins and Dr. Kathryn Walder for antibodies and assistance with Ankyrin analysis. We thank Dr. Young Truong (Department of Biostatistics at UNC) for statistical help. Dr. Damaris Lorenzo provided assistance with mouse breeding.

